# Voltage-dependent volume regulation controls epithelial cell extrusion and morphology

**DOI:** 10.1101/2023.03.13.532421

**Authors:** Saranne J. Mitchell, Carlos Pardo-Pastor, Thomas A. Zangle, Jody Rosenblatt

## Abstract

Epithelial cells work collectively to provide a protective barrier, yet also turn over rapidly by cell death and division. If the number of dying cells does not match those dividing, the barrier would vanish, or tumors can form. Mechanical forces and the <u>s</u>tretch-<u>a</u>ctivated ion <u>c</u>hannel (SAC) Piezo1 link both processes; stretch promotes cell division and crowding triggers cell death by initiating live cell extrusion^1,2^. However, it was not clear how particular cells within a crowded region are selected for extrusion. Here, we show that individual cells transiently shrink via water loss before they extrude. Artificially inducing cell shrinkage by increasing extracellular osmolarity is sufficient to induce cell extrusion. Pre-extrusion cell shrinkage requires the voltage-gated potassium channels Kv1.1 and Kv1.2 and the chloride channel SWELL1, upstream of Piezo1. Activation of these voltage-gated channels requires the mechano-sensitive Epithelial Sodium Channel, ENaC, acting as the earliest crowd-sensing step. Imaging with a voltage dye indicated that epithelial cells lose membrane potential as they become crowded and smaller, yet those selected for extrusion are markedly more depolarized than their neighbours. Loss of any of these channels in crowded conditions causes epithelial buckling, highlighting an important role for voltage and water regulation in controlling epithelial shape as well as extrusion. Thus, ENaC causes cells with similar membrane potentials to slowly shrink with compression but those with reduced membrane potentials to be eliminated by extrusion, suggesting a chief driver of cell death stems from insufficient energy to maintain cell membrane potential.

## MAIN

Cell extrusion promotes cell death to maintain homeostatic epithelial cell numbers, using a mechanism conserved from sea sponge to humans. During extrusion, a live cell is ejected apically by coordinated basolateral actomyosin contraction of the extruding cell and its neighbors^1^. After extrusion, the cell dies due to lack of survival signaling. We previously discovered that Piezo1 activates <u>l</u>ive <u>c</u>ell <u>e</u>xtrusion (LCE) in response to crowding to maintain constant epithelial cell numbers^1,2^. Crowding activation of Piezo1 triggers a canonical pathway that also extrudes cells triggered to die by apoptosis. To extrude, cells produce the lipid Sphingosine 1-Phosphate (S1P), which binds the G-protein-coupled receptor S1P_2_ to activate Rho-mediated actomyosin contraction needed for extrusion^2–4^. As crowding-induced cell extrusion drives most epithelia cell death, an important outstanding question is which cells within a crowded epithelial field are chosen for extrusion? In sparse epithelia, topological defects within a normal hexagonal-packed epithelial promote outlier cells to extrude^5,6^. Additionally, *C. elegans* and mammalian cells with DNA damage or replicative stress are selected for extrusion^6,7^. However, as most cell extrusions occur in crowded regions, where cells do not divide or have obvious topological defects, what marks most cells for extrusion has remained a mystery.

One possible model is Darwinian, where cells that are fitter remain in epithelia while weaker ones are marked for extrusion and death^8^. One possible measure of cell fitness in this model is cell mass. Cells with higher dry cell mass might be fitter in terms of protein and macromolecule production and maintenance. To test this possibility, we adapted <u>q</u>uantitative <u>p</u>hase <u>i</u>maging (QPI) to analyze dry mass changes over time in Madin-Darby Canine Kidney II (MDCKII) epithelial monolayers (see methods)^9,10^. However, QPI revealed that dry mass does not change prior to cell extrusion, unlike the clear mass increase before a cell divides (Supplemental Fig.1A-B and video 1, representing 31 cells from 12 total cell islands), disfavoring cell mass change as a factor for selecting LCE.

**Supplementary Figure 1.**
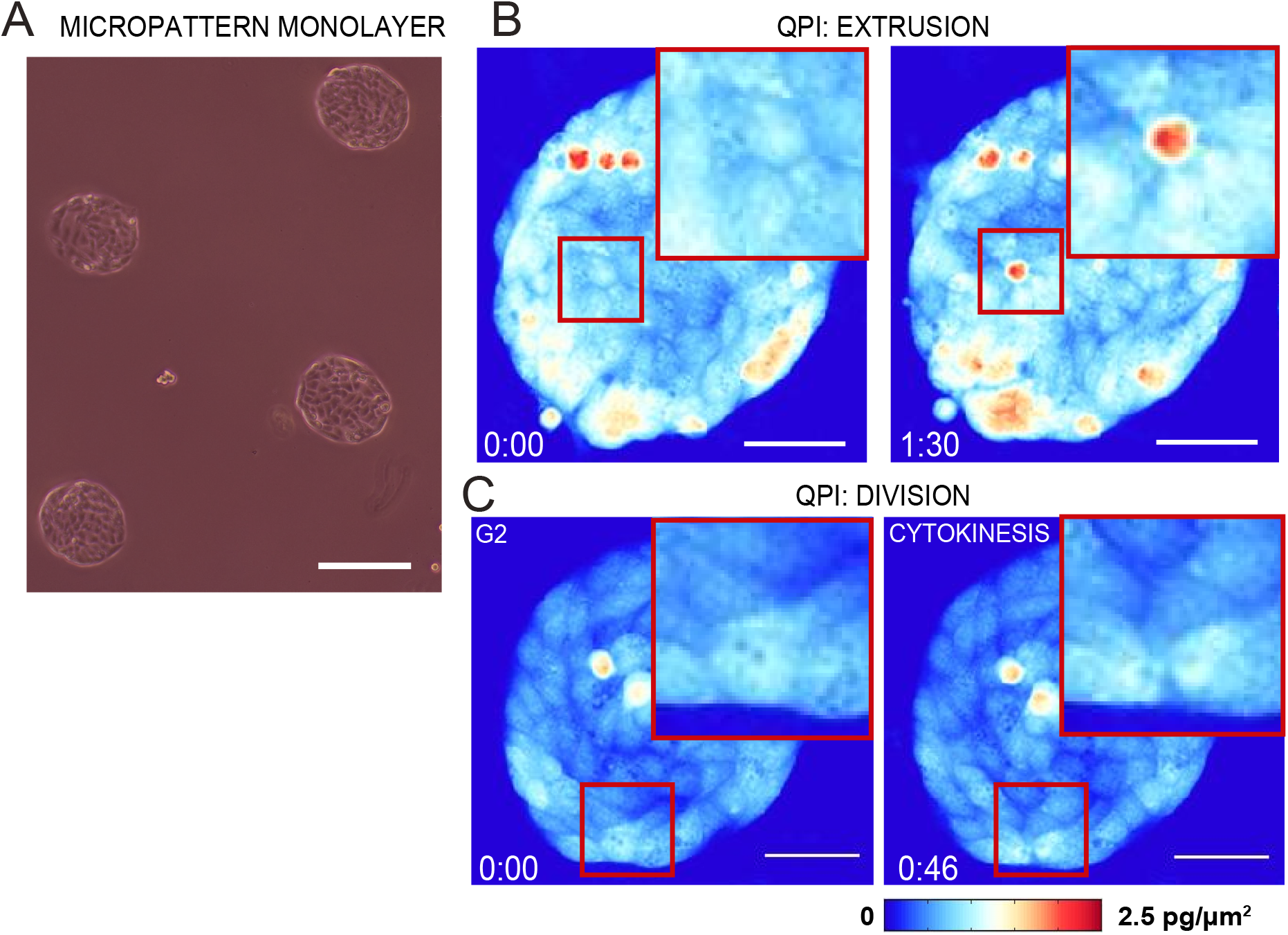
Quantitative phase imaging (QPI) shows unaltered dry mass in cells that will extrude. **A,** As QPI requires a reference area without cells throughout imaging, we grew MDCKII monolayers on confined 100μm islands within a 35 mm dish by plasma treating patterned areas, as in this example. Scale bar=100μm. **B,** QPI stills from Supplemental video 1, representing 31 other cells from 12 QPI islands that similarly show dry mass before cell extrusion, where blue is low cell mass and red is higher mass. Note mass increases after cell extrusion, as the density gets higher with cell contraction (h:mm). **C,** By contrast, another cell, in red box, increases mass before dividing. Scale bar = 50μm.

While investigating cell mass, we noted a consistent, transient increase in cell-cell junction brightness by phase microscopy that lasts ~6.5 minutes before a cell extrudes (Fig.1A, B, and Supplemental Video 2). To readily assay the transient phase brightness, we developed the ‘lightning assay’ that relies on intensity thresholding of junctional brightness surrounding cells before they extrude (Fig.1C&D). By contrast, less than 0.03% of cells filmed (8 in 750) exhibit junctional brightness without extruding (Fig. 1E). Increased phase brightness might indicate a decrease in cell volume or shape. To test if transient volume loss accounts for the phase-brightness before extrusion, we expressed cytoplasmic GFP mosaically in MDCKII cells and measured cell volume throughout extrusion by confocal timelapse microscopy. We find that live cells experience a transient area and volume loss before extruding that corresponds to the junctional brightness measured in our lightning assay (Fig. 1D, F&G). Given this direct correlation between phase brightness and cell volume, we used the lightning assay for measuring Homeostatic Early Shrinkage (HES) before LCE. Importantly, we find that HES before extrusion occurs in only crowded regions, as measured by cell density; the few cells that shrink without extruding do so in uncrowded epithelial regions (Fig. 1H). Thus, crowding followed by single cell shrinkage appear to initiate cell extrusion.

**Figure 1:**
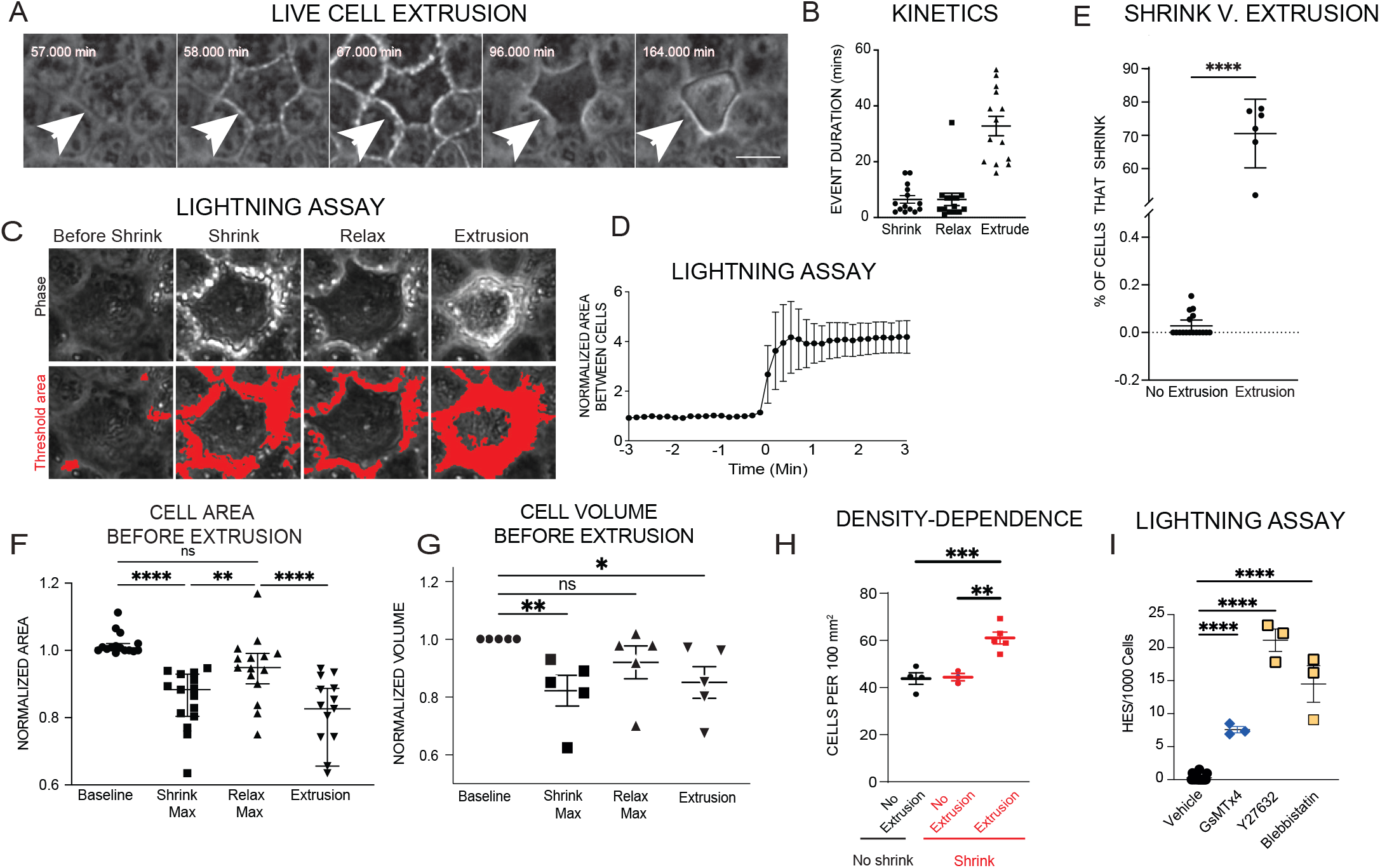
Live cells shrink briefly before extruding in dense regions. **A,** Stills from a phase timelapse movie show junctional lightning (white arrowhead) around a cell before it extrudes. Scale bar=10μm **B,** Kinetics of live cell extrusion denoting average time (in minutes) cells experience shrink, relax, and from shrink to extrusion, extrusion, mean ± SEM. **C,** Lightning assay, showing representative stills from time-lapse phase microscopy where white space is thresholded (red, bottom) at key points during extrusion, quantified in (**D)** as mean ± SEM of normalized white area around cell, where t0 is shrink; n=10 cells. **E,** Percent cells that shrink before extruding ± SEM and P from unpaired T-test using n=10 biological replicates. **F**, Normalized area mean ± SEM of cells at each defined step: baseline, shrink maximum, relax maximum and extrusion; n=15 cells; p values from two-way ANOVA with Dunnett’s multiple comparisons test. **G,** Normalized mean volume ± SEM of cells at baseline, shrink maximum, relax maximum, and onset of extrusion; n=5 cells with p values from two-way ANOVA with Dunnett’s multiple comparisons test. **H,** Regional density-dependence ofcell shrinkage (red) with or without extrusion, compared to no shrink (black); as mean ± SEM, n=3 with one-way ANOVA with Dunnett’s multiple comparisons test. **I,** Mean rate of HES with inhibitors of cell extrusion ± SEM; n=3, using one way ANOVA with Dunnett’s multiple comparisons test. For all graphs, ****P<0.0001, ***P<0.001, **P<0.01 or *P=0.0124.

**Supplemental Figure 2.**
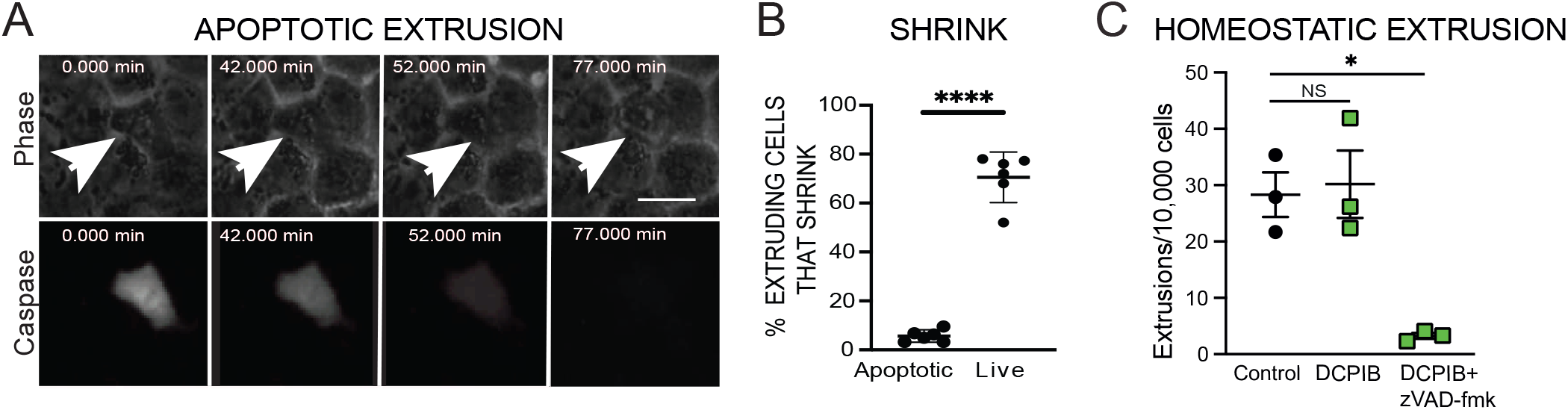
HES does not occur before apoptotic cell extrusion. **A,** Stills from a phase (top) timelapse and fluorescent apoptosis (Cell Event-caspase) marker (bottom) before apoptotic extrusion. Scale=1Oμm. **B,** Mean percentage of cells ± SEM that shrink by lightning assay before extruding. n=5; ****P<0.0001 by an unpaired T-test. **C,** Mean extrusion rate ± SEM in monolayers treated with SWELL1 inhibitor ±zVAD-FMK; n=3; where *P=0.0104 values are from one-way ANOVA with Sidak’s multiple comparisons test.

While ~70% of extruding cells undergo HES, it was not clear why the remaining 30% do not shrink. One possibility is that this minor population of non-shrinking extrusions were apoptotic, which accounts for ~20-30% of extruding cells at steady state in cultured epithelia^1^. Combining a fluorescent reporter of the cleaved caspase-3 apoptotic marker with the lightning assay, we find cell shrinkage rarely occurs (~3%) before apoptotic extrusion (Supplemental Fig 2A-B, Supplemental Video 3). Thus, HES typically precedes LCE but not apoptotic extrusion.

Transient cell volume loss could be due to myosin II contraction, which occurs before apoptotic extrusion^11,12^. To investigate if myosin contraction is important for HES, we inhibited ROCK or myosin II and measured cell shrinkage with our lightning assay. We found blocking myosin II contraction increased the number of cells shrinking by ~23X compared to untreated controls (Fig. 1I), suggesting HES occurs independently from myosin contractility. Additionally, HES did not cause blebbing, typically associated with myosin II contraction^13,14^, further ruling out a role for myosin contraction in HES. As HES occurs in crowded regions, we investigated if Piezo1, the SAC required for crowding-induced live cell extrusion^2^, might instead control transient cell volume loss. However, we find that the SAC inhibitor GsMTx4 does not block HES, suggesting Piezo1 acts downstream of HES (Fig. 1I). This was surprising, as our previous studies suggested Piezo1 as the earliest mechano-sensor to trigger extrusion in response to crowding^1^.

Having ruled out dry mass, contractility, and early extrusion regulators as controllers of HES, we next investigated if water efflux may regulate the transient volume loss cells experience before extrusion. Cellular water regulation is critical for controlled epithelial cell death^7,15,16^ and could account for the volume changes seen before LCE. To first test if water loss is sufficient to initiate LCE, we incubated MDCKII monolayers in media containing increasing osmolarities (hypotonic to hypertonic) for ten minutes before returning to isotonic, control medium to mimic the duration of shrinkage before extrusion (Supplemental Video 4&5, Fig. 2A). While hypotonic medium had no impact on cell extrusion (Fig. 2A), 15-20% hypertonicity dramatically increased both cell shrinkage and LCE rates (Fig. 2A&B), with higher concentrations causing so much extrusion that it destroyed the monolayer (data not shown). Although increasing hypertonicity can provoke <u>A</u>poptotic <u>V</u>olume <u>D</u>ecrease (AVD)^12,17^, the lack of fluorescence associated with cleaved caspase-3/7 apoptotic cell probe indicated that hypertonic treatment induces only LCE (Fig. 2B). Further, the pan caspase inhibitor zVAD-fmk does not prevent 20% hypertonic shrink-induced extrusions (Supplemental Fig. 3A&BB), ruling out AVD as a mechanism for hypertonic-induced extrusion. These findings further support that HES controls live but not apoptotic cell extrusion. Interestingly, although the entire monolayer was treated with 20% hypertonic solution, not all cells shrunk equally. Only cells that shrank 20±3% as measured by cytoplasmic GFP volume extrude, whereas those shrinking 11±2.5% do not (Fig. C2C). Thus, experimentally reducing cell volume ~20% is sufficient to trigger individual cells to extrude, which we refer to as <u>o</u>smotic-<u>i</u>nduced <u>c</u>ell <u>e</u>xtrusion (OICE).

**Figure 2:**
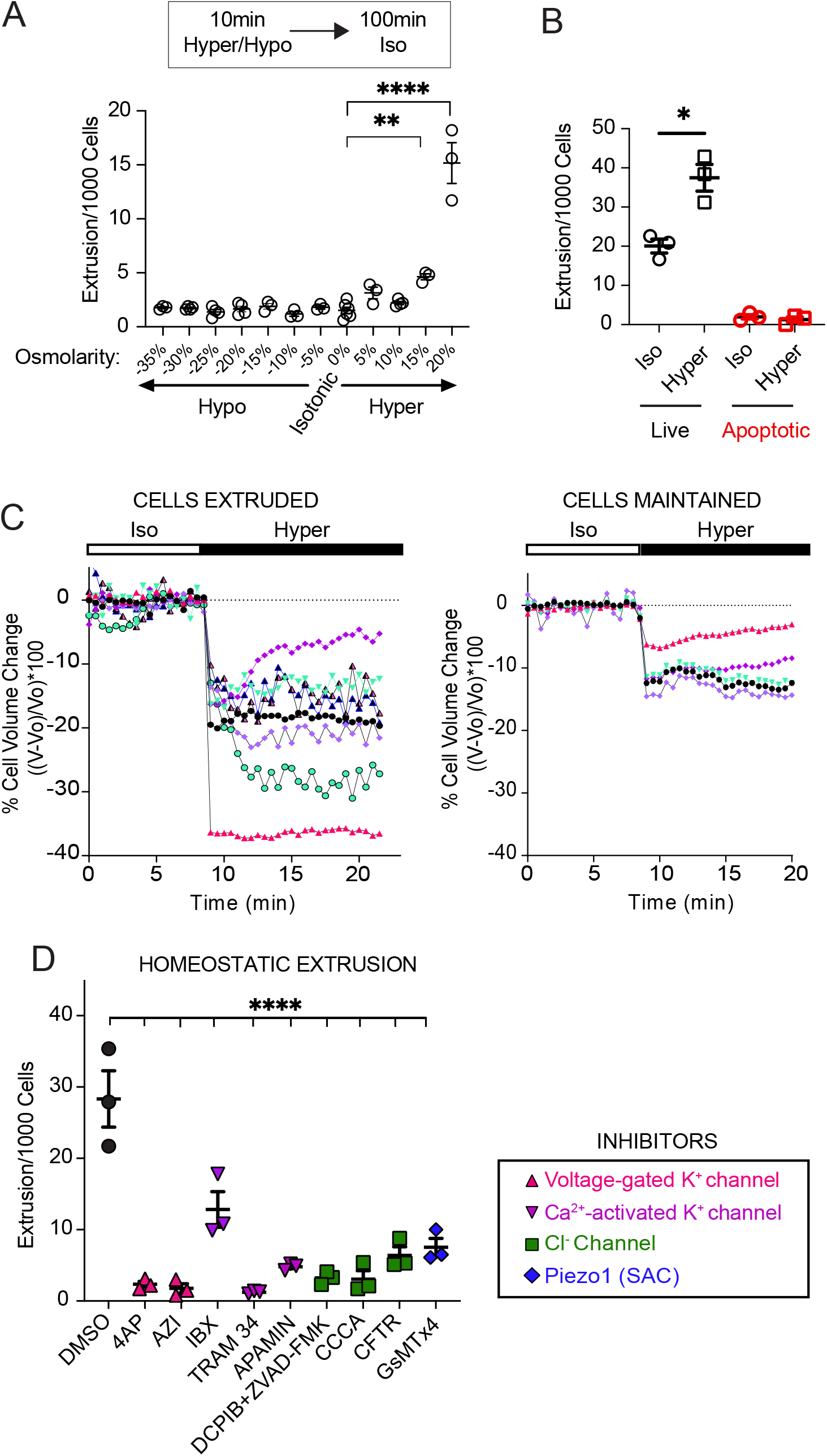
Water loss is sufficient to induce live cell extrusion. **A,** Mean MDCKII extrusion rate ± SEM following incubation in media with increasing osmolarity for 100’, scored by immunostaining for actin and DNA, with n=3 experiments; where **P= 0.0068; ****P<0.0001 by one way ANOVA with Dunnett’s multiple comparisons test. **B,** Mean extrusion rates ±SEM in normal medium or 100’ following 10 minute 20% hypertonic shock, where apoptotic extrusion is scored by immunostaining with cleaved caspase-3 antibody; n=3 experiments; and *P=0.0105 is from an unpaired T-test. **C,** Volume changes (measured by confocal sectioning of mosaically expressed GFP) over time in cells shifted from isotonic to 20% hypertonic media that extrude (left) or do not (right), n=5. **D,** Number of extrusions in MDCKII monolayers incubated with volume regulating inhibitors and scored by immunostaining for actin and DNA; n=3; where****P<0.0001 by one way ANOVA with Dun-nett’s multiple comparisons test. Inhibitor key describes volume regulating channel family inhibitor target by assigned color and icon.

**Supplemental Figure 3.**
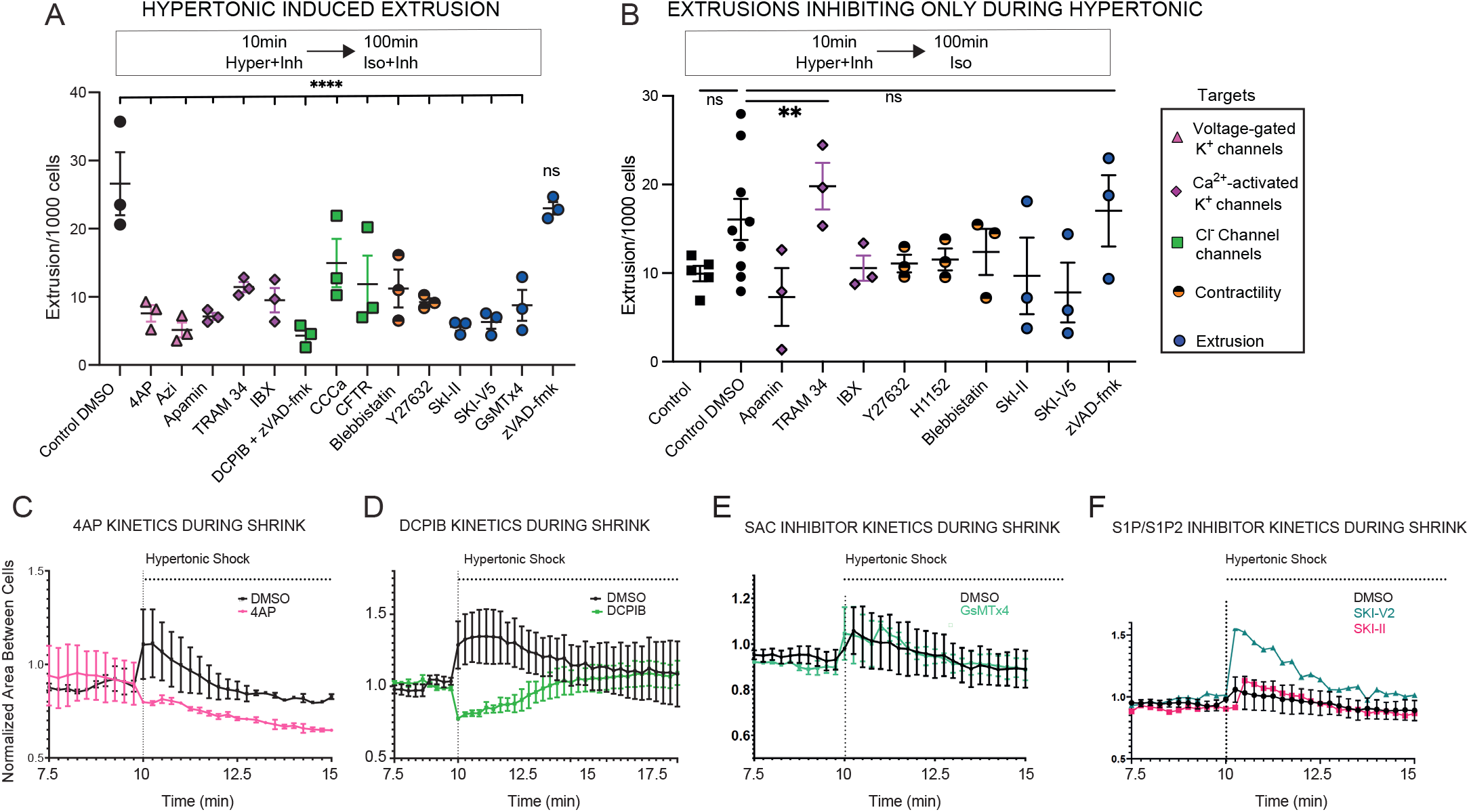
Ion channel inhibitors impact on extrusion and shrink. **A,** Mean extrusion rate ± SEM in MDCKII monolayers treated with inhibitors before, during, and after hypertonic shock; n=3; where ****P=0.0001 by one way ANOVA with Dunnett’s multiple comparisons test. **B,** Mean extrusion rate ± SEM in monolayers treated with inhibitors only during hypertonic shock. n=3 experiments where **P=0.0024 is from two-way ANOVA with Dunnett’s multiple comparisons test. Inhibitor key for A and B describes ion channel family inhibitor target by color icon. Representative “Lightning assays” where increased space around the cells indicates cell shrinkage in the presences of: 4AP (KV1.1&KV1.2 inhibitor) (**C**), DCPIB (SWELL1 inhibitor) (**D**), SAC inhibitor GsMTx4 (**E**), or S1P/S1P_2_ inhibitors (**F**), compared to DMSO controls before and during hypertonic media incubation (mins).

Since potassium (K^+^) and chloride (Cl^-^) efflux drive water loss during AVD^23^, we investigated if they control HES and extrusion. Using a panel of K^+^ and Cl^-^ channel inhibitors, we assayed for those that block homeostatic extrusion rates by timelapse microscopy and immunostained samples. The inhibitor carrier, dimethyl sulfoxide (DMSO), can change water permeability of cells^18^, however it did not decrease the basal rate of extrusion, ruling out nonspecific effects (Fig. 2D and A3A). Although most inhibitors were not toxic, the Cl^-^ channel SWELL1 is essential for cell survival^19^, requiring the addition of caspase inhibitor zVAD-fmk to its inhibitor DCPIB for all our assays (Fig 2D, Supplementary Fig 2C). Remarkably, all inhibitors tested significantly reduced extrusion rates during normal homeostatic turnover to levels seen when SAC channels are inhibited with GSMTx4 (Fig. 2D and Supplementary Fig 3A). Interestingly, 4-AP, an inhibitor of voltage-gated Kv1.1 and Kv1.2 channels, also blocks apoptotic extrusion^20^, suggesting a conserved activator for all cell extrusions (Fig 2D, Supplementary Fig 3A). Other channels are implicated in pleotropic diseases, like Kv11.1 in carcinomas^21,22^ and heart arrhythmias^23^ and the cystic fibrosis transmembrane receptor (CFTR) in cystic fibrosis^24^, suggesting possible new roles for misregulated extrusion in these diseases. While these findings were surprising and suggest interesting future studies, here, we focus solely on those that control the HES step.

To determine which K^+^ and Cl^-^ channels, if any, were needed for HES, we first tested which inhibitors blocked extrusions when present only during the 10-minute hypertonic treatment of OICE. Timelapse phase microscopy revealed that only early inhibition of SACs, Kv channels, and Cl^-^ channels stopped extrusion (Fig. 3A and Supplementary Fig. 3B). Even though GsMTx4 does not stop cell shrinkage, it inhibits cell extrusion when inhibited only in the first 10-minutes, signifying its role early during extrusion pathway, yet downstream of cell shrinkage (Fig. 1I, 2D). Thus, we next investigated if the voltage-gated K^+^ and Cl^-^ channels required in early stages of extrusion regulate both osmotic-induced and homeostatic shrinkage before extrusion.

**Figure 3:**
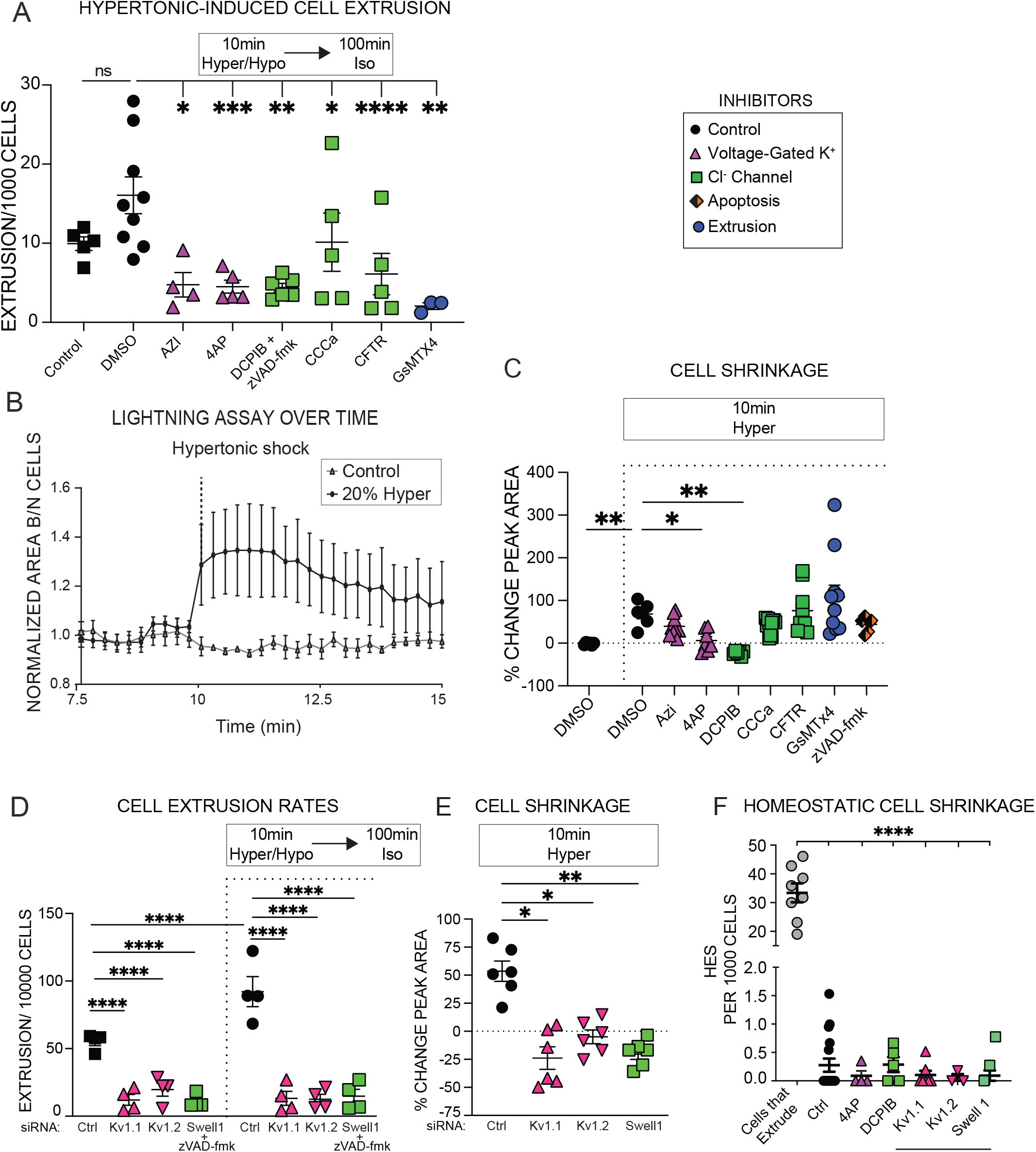
Homeostatic live cell extrusion requires K^+^ voltage-gated and Cl^-^ channel activation. **A,** Mean extrusion rate ± SEM from monolayers treated with inhibitors only during the 10’ 20% hypertonic treatment n=3;P values from twoway ANOVA with Dunnett’s multiple comparisons test. **B,** Adapted lightning assay for a crowded epithelial field following hypertonic treatment, quantified over time. **C,** Mean percent peak change in area surrounding cells (from adapted lightning assay) ± SEM with inhibitors during 20% hypertonic challenge, n=6; with P values from a two-way ANOVA with Dunnett’s multiple comparisons test. Channel nhibitor key (for B & C). **D,** Mean extrusion rate ± SEM of MDCKIIs with siRNA-knockdowns ± 20% hypertonic challenge, compared to controls; n=4; P values from two-way ANOVA with Dunnett’s multiple comparisons test. **E,** Percent change of peak area (from adapted lightning assay) ± SEM during hypertonic challenge of siRNA-knockdowns, compared to controls, n=6; with P values from one-way ANOVA with Dunnett’s multiple comparisons test. **F,** Mean rate of HES ± SEM when treated with 4AP, DCPIB, or with Kv1.1, Kv1.2 or SWELL1 siRNA knockdowns, compared to control background cells or those that shrink before extruding; n=3; where P values are from two-way ANOVA with Dunnett’s multiple comparisons test. For all graphs, ****P<0.0001, ***P<0.001, **P<0.0100, *P<0.05

To identify which ion channels regulate HES, we first adapted our lightning assay (Fig 1D&E) to measure increases in light intensity of all cell-cell junctions within a crowded field in response to a 20% hypertonic shock, (Fig. 3B, normalized to isotonic medium), as a proxy for cell volume loss. This assay identified only DCPIB (combined with zVAD-fmk) and 4-AP, which target the chloride channel SWELL 1 and the voltage-gated channels Kv1.1, Kv1.2, respectively, as inhibitors of both hypertonic-induced cell shrinkage and extrusion (Fig. 3A&C & Supplementary Fig. 3A-DD). Immunostaining showed that all three channels localize to the cell apex, with Kv1.1 co-localizing with ZO-1 at tight and tri-cellular junctions, regions known to act as tension sensors^25^ (Supplementary Fig. 4A). We used siRNA-mediated knockdown to test the roles of Kv1.1, Kv1.2, and SWELL 1, the targets of 4-AP and DCPIB, in both OICE and HES (Suppl. Fig. 4B&C). We found that knockdown of any of these channels prevented HES and OICE (Fig. 3D-F), suggesting that they regulate both cell shrinkage and extrusion during normal cell turnover.

The fact that Kv1.1, Kv1.2 are voltage-gated channels suggested that membrane depolarization could trigger both HES and LCE^26^. Although most frequently studied in neuronal cells, electrical membrane potentials are critical for all cells, requiring between 30-70% of all the energy consumed within a cell, depending on the cell type^27^. Na^+^/K^+^ ATPases maintain polarized ions with intracellular K^+^ and Cl^-^ and extracellular Na^+^ and organic anions to create these membrane potentials. While individual cells create membrane potentials, epithelial cells tethered together with tight, adherens, and gap junctions, work collectively to maintain a transepithelial potential difference of between 1-46mV, with a net negative charge on the apical side^28,29^. Interestingly, imposing a direct current that reinforces this polarity (from basal to apical) enhances and even repairs impaired epithelial cell-cell junctions, whereas reversing the current (from apical to basal) causes cells to contract and detach from each other^30^, reminiscent of the cell shrinkage and junctional lightning between cells before extrusion or with hypertonic treatment (Figs. 1F&3C).

Since voltage-gated channels control cell shrinkage and experimentally depolarizing a whole epithelial monolayer produces similar shrinkage of all cells, we next investigated if cells depolarize their membranes before they shrink. To visualize membrane potential, we filmed MDCKII monolayers with the slow-response potential-sensitive probe DiBAC3(4), which increases fluorescence with membrane depolarization31-33. Unlike excitable tissues that rapidly depolarize in an all or nothing reversible manner, we found that MDCKII cells within the monolayer have different membrane potentials; only some cells become DiBAC3(4)-brighter, depolarizing as they crowd and maintaining this depolarized state (Fig. 4A&B). This shows that despite epithelial cells being connected with gap junctions, they are not fully electrically coupled. This also supports our previous finding that cells become smaller in regions of crowding1,34 and suggests that individual cell volume/size may be governed by existing Na+-K+ pumps or ATP available to maintain resting membrane potential35. Importantly, we found that before cells shrink and extrude, they depolarize significantly, becoming DiBAC3(4)-brighter (Fig. 4C-E, Supplemental Video 7), whereas cells that do not shrink, do not depolarize, linking shrinkage to depolarization (Fig. 4E&F). Strikingly, most cells that depolarize before shrinking and extruding have markedly more positive membrane potentials than the cells directly neighboring them (Fig. 4D and Supplemental Video 6). Thus, as cells within a monolayer experience crowding, those with similar membrane potentials remain in the field and shrink slightly whereas those with strikingly lower potentials are eliminated by extrusion. From our measurements of thresholds for shrinkage, the difference in whether a cell extrudes or not may be due to whether the cell shrinks enough (20±3%) to elicit extrusion.

**Figure 4:**
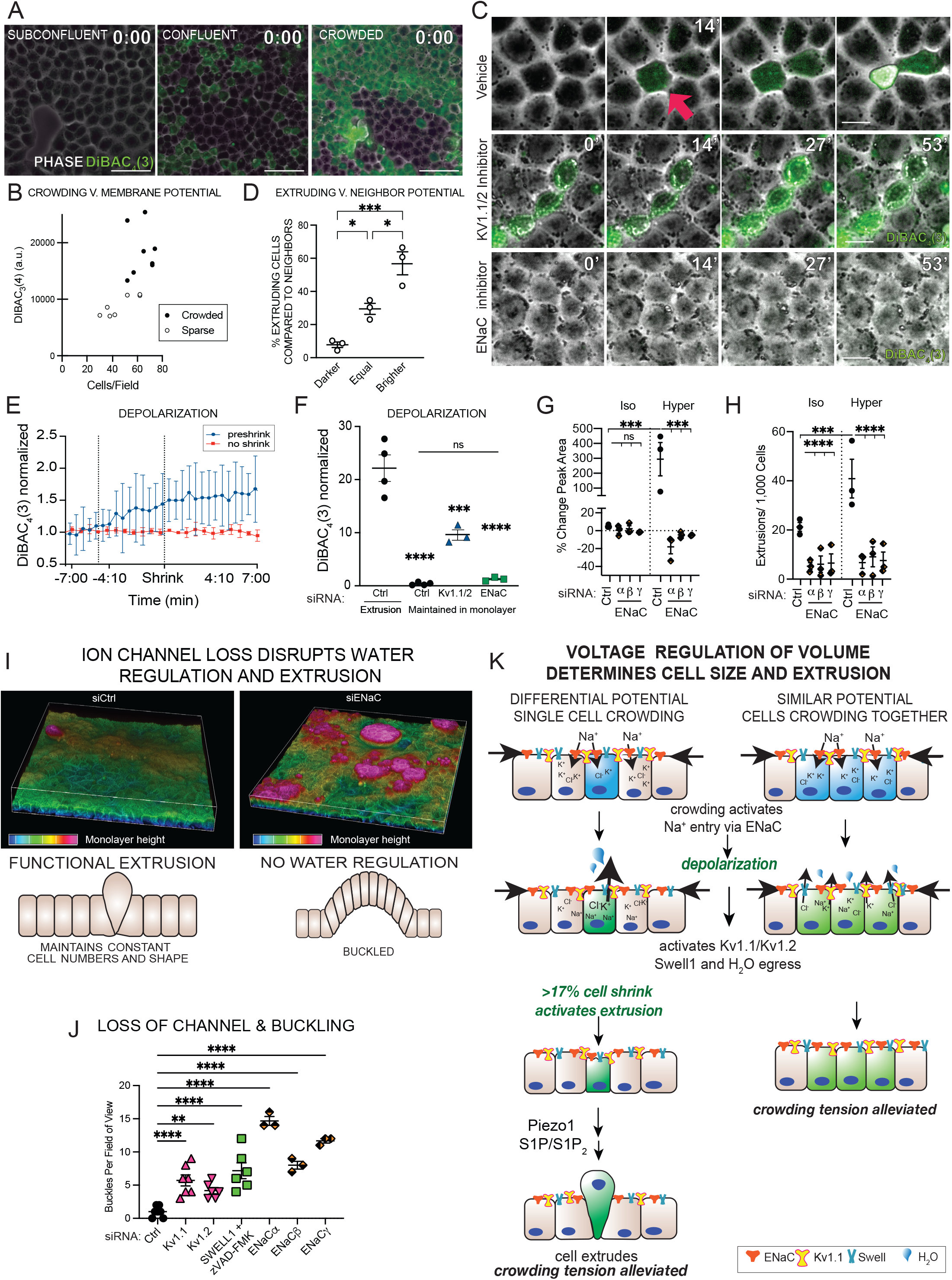
ENaC-dependent depolarization activates cellular shrinkage and extrusion. **A,** Membrane depolarization, measured by DiBAC_3_(4) fluorescence (greater depolarization with higher fluorescence), in sub-confluent, confluent, and crowded MDCKII monolayers. Scale bar =50μm. **B,** Density-dependence of membrane potential, as measured by DiBAC_3_(4) fluorescence, n=3. **C,** Timelapse (mins) measuring membrane depolarization before homeostatic extrusion (top panel, red arrow indicating a cell before extrusion), in the presence of 4AP (KV1.1/2 inhibitor, middle panel), and with amiloride (ENaC inhibitor, bottom panel). Scale bars =10μm. **D,** Mean cells ± SEM with darker, equal, or brighter DiBAC_3_(4) staining, compared to surrounding neighbouring cells before extruding; n= 3 experiments; P values from one-way ANOVA with Tukey’s multiple comparisons test. **E,** Normalized DiBAC_3_(4) fluorescence, as mean ± SEM from cells that undergo HES/extrusion (blue) or not (red) over time, n=6. **F,** Mean membrane polarization status ± SEM, as measured by DiBAC_3_(4) fluorescence of cells before they extrude compared to non-extruding cells in background or when Kv1.1/Kv1.2 or ENaC are inhibited with 4AP or amiloride; n=3 experiments; P values from one-way ANOVA with Tukey’s multiple comparisons test. **G,** Peak mean percent area change ± SEM (by adapted lightning assay) of siRNA-mediated ENaC-α, β, or γ knockdown in MDCKs with and without 20% hypertonic challenge, compared to non-targeted controls, n=3 for each siRNA, with P values from a two-way ANOVA with Dunnett’s multiple comparisons test. **H,** Mean extrusion rates ± SEM of ENaC-α, β, or γ siRNA-mediated knockdown with and without 20% hypertonic challenge, compared to control, n=3; with P values from two-way ANOVA with Dunnett’s multiple comparisons test. **I,** Representative volume view of control and ion channel knocked down monolayers, with alpha color bit coded bar indicating depth 0.0-16.0μm (blue-pink). Schematics below depict how functional extrusion maintains monolayer shape whereas disrupting water regulation leads to buckling. **J,** Mean number of epithelial buckles per field ± SEM 48 hours post siRNA-treatment compared to non-target controls; n=3; with P values from two-way ANOVA with Dunnett’s multiple comparisons test. **K,** Model for how ENaC/Kv1.1/Kv1.2/Swell1-dependent water regulation controls cell size and extrusion, depending on cellular membrane potential. ***Left:*** If one cell is more depolarized than its neighboring cells, crowd activation of ENaC will preferentially activate voltage-gated potassium channels in this cell, causing shrink. II this shrink reaches a threshold of ~20%, it will activate the extrusion pathway. Cell elimination by extrusion will alleviate crowding pressure in the monolayer and maintain constant densities. ***Right:*** Crowd activation of ENaC in cells with equivalent membrane potentials cause moderate voltage-gated potassium-dependent water loss across all cells, resulting in slight cell shrinkage but not enough to activate cell extrusion. Here, slight shrinkage also helps accommodate overall crowding of the monolayer. In this way, crowding can set up competition to eliminate cells with the lowest energy that cannot maintain a sufficient membrane potential, a significant energy demand for the cell. For all graphs, ****P<0.0001, ***P<0.001, **P<0.0100.

**S. Figure 4.**
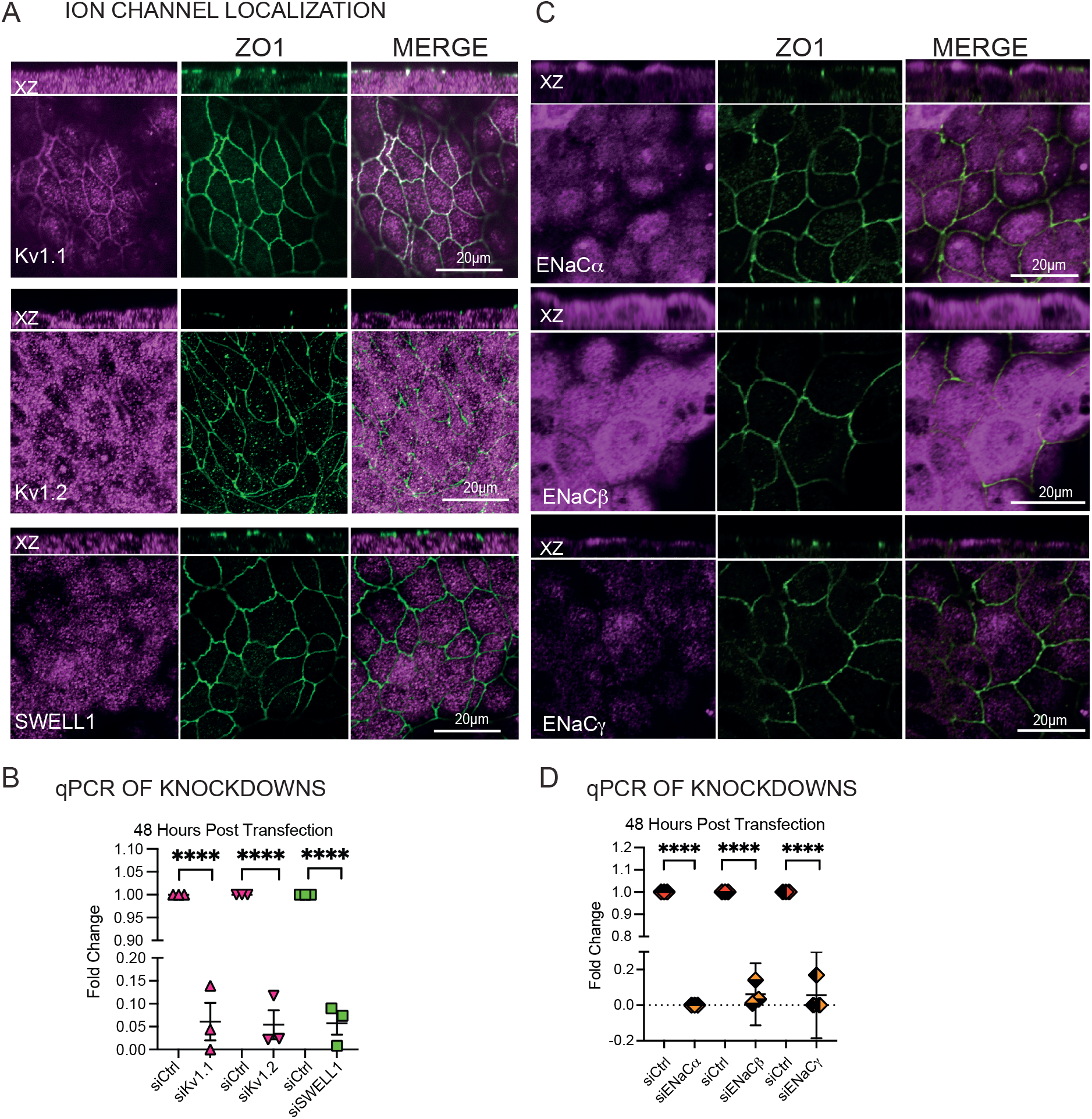
HES required ion chanel location and siRNA knock down validation. **A,** Confocal representative projections and xz images of Kv1.1/1.2, SWELL1 or ENaC α, β, or γ (magenta) with apical tricellular junction protein ZO-1 (green). XY Scale bar = 20μm. **B**, Table of primer sequences used to quantify siRNA-mediated knockdown for each channel and control GAPDH by qPCR. **C**, Scatter plots showing mRNA fold change (2^ΔΔCt^) at 48 hours post transfection; n=3; where ****P=0.0001 from an unpaired T-Test.

**Supplemental Figure 5.**
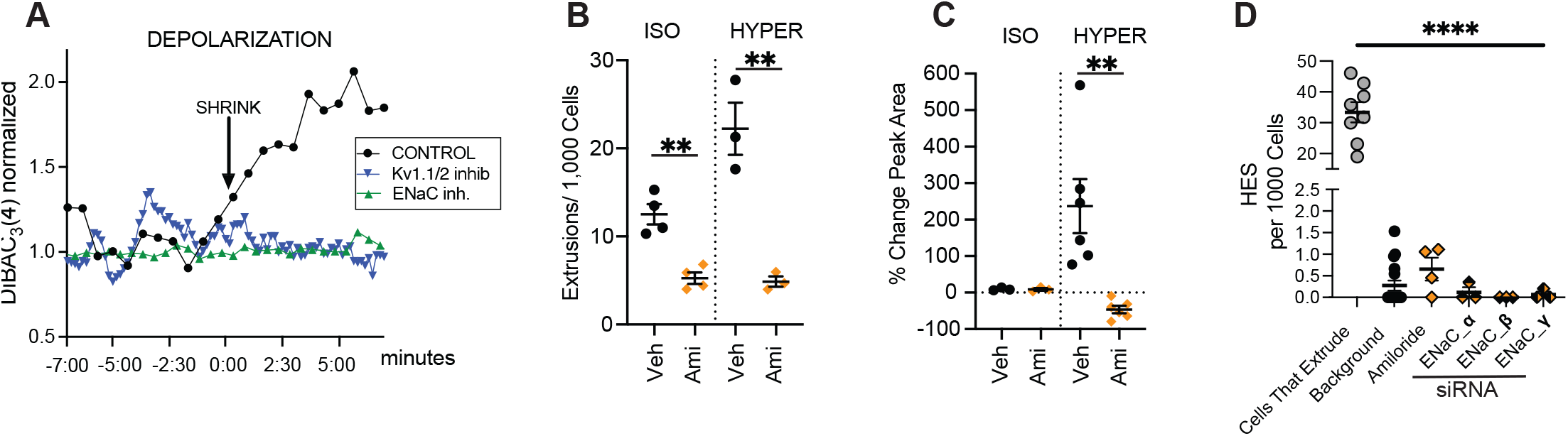
Inhibition of ENaC on membrane potential and extrusion. **A,** Mean extrusion rate ± SEM of MDCKIIs treated with amiloride, compared to DMSO controls. n=4 homeostatic (isotonic) and n=3 for hypertonic challenges, where **P<0.0100 by an unpaired t-test. **B,** Mean percent peak change in area ± SEM of cells (by lightning assay) with amiloride during isotonic or 20% hypertonic challenge, compared to DMSO controls, n=3 for homeostatic (isotonic) and n=6 for hypertonic challenge, where **P=0.0036 derives from an unpaired t-test. **C,** Representative depolarization of MDCKII cell over time (mm:ss) treated with DMSO control, Kv1.1/1.2 inhibitor 4AP, or ENaC inhibitor amiloride. **D,** Mean rate of HES ± SEM (by lightning assay) in non-extruding cells when treated with amiloride, or with ENaC-a, β or γ siRNA, compared to cells that shrink before extruding; n=3 where ****P<0.0001 is derived from two-way ANOVA with Dunnett’s multiple comparisons test.

Membrane depolarization before voltage-gated dependent cell shrinkage suggests that another channel activates Kv1.1 and Kv1.2. One attractive candidate is the Epithelial Sodium Channel (ENaC)—a highly conserved, mechanically activated, apically localized (Supplementary Fig. 4D) ion channel that causes membrane depolarization via Na^+^ entry^36,37^. Inhibiting ENaC with amiloride or knocking down any of its subunits (a, β, or γ, Supplementary Figs. 4 B&C) prevented both HES and OICE (Fig. 4G&H). Importantly, ENaC inhibition with amiloride also prevented depolarization of all epithelial cells in response to hypertonic challenge or during steady state turnover (Fig. 4C&F). Because inhibiting Kv1.1 and Kv1.2 with 4-AP does not prevent membrane depolarization (Fig. 4C&F and Supplementary Fig. 5C), we surmise that ENaC triggers membrane depolarization, which, in turn, activates the voltage-gated potassium channels Kv1.1 and Kv1.2.

If ENaC, Kv1.1, Kv1.2, or SWELL1 control epithelial cell shrinkage and extrusion, their loss should change epithelial morphology. We find that deregulating water and extrusion from knockdown of any of these channels, in fact, causes epithelial buckling (Fig 4 I&J). Compared to control monolayers that shrink and extrude in crowded areas^34^ (Fig1.H), those that lack any one of these channels do not undergo HES or OICE (Figs. 3D-F and Supplementary Figs. 5A, B, & D) but instead form progressive buckles as cells crowd (Supplemental Video 8). Thus, cell shrinkage and extrusion alleviate crowding of epithelia sheets to maintain constant numbers and morphology; when epithelia cannot regulate their volume, they can become buckled (see schematic below Fig. 4K). Presumably, in wild type tissues, cells must decrease their plasma membranes by endocytosis or other mechanisms to become smaller overall. Decreasing only cell volume without surface area may instead lead to cell pseudo-stratification with crowding. Thus, we suggest that water movement and pressure is sufficient to control epithelial cell numbers and morphologies. Potentially, the set points for water regulation in different epithelia could govern whether tissues become buckled like the gut, kidney, and lungs^38,39^ or pseudostratified, like neuroepithelia^40^ and some adenomas^41,42^, reinforcing the role of hydrodynamics in morphogenesis^43^

While several models could explain our findings, we favor the following model for how crowding, Na^+^, K^+^, and Cl^-^ control epithelial cell volume and turnover (Fig. 4K). As epithelial cells divide and accumulate, they cause crowding within a layer, which activates the tension sensitive sodium channel ENaC that depolarizes cells. Membrane depolarization by ENaC activates Kv1.1 and Kv1.2 that with Swell1 drive water egress and cell shrinkage. Cells with similar membrane potentials will experience depolarization but only slight water loss and shrinkage within a crowded field. However, ENaC-dependent depolarization in a cell that is already more depolarized than its neighbors will reach the threshold for Kv activation more readily. Moreover, mechanical forces can reduce the voltage needed to activate Kv channels and increase their maximal current^44,45^. Considering this, ENaC-Kv coupling will be more efficient in a crowded, partially depolarized cell than in its neighbours, further amplifying the initial difference. Thus, a cell under mechanical strain would shrink more readily and significantly, with those shrinking >17% of their original volumes triggering extrusion. Cell extrusion would also serve to alleviate crowding and ENaC activation in nearby cells, inhibiting the axis and preventing excessive extrusion. Additionally, the differential in membrane potentials between a cell targeted for extrusion and its neighbors could set up a directional current, which has been shown to activate migration during wound healing^46^.

While our study revealed a new role for water regulation and membrane potential in regulating epithelial cell extrusion and morphology, it also opened several new questions for further investigation. For instance, shrinkage seems to amplify the crowding experienced in a cell that will extrude, but we still lack information as to how transient cell shrinkage activates Piezo1 to trigger S1P signaling vesicles, essential to extrusion^2–4^. Further, while we identified potassium and chloride channels essential for HES, it is likely that aquaporins may work in conjunction with these channels to coordinate water loss; identifying which will be a future goal. Our findings also revealed the importance of other ion channels in regulating extrusion and it will be important to define what roles they play in controlling extrusion. For example, malfunction of ENaC and CFTR are known drivers of Cystic Fibrosis^24,47,48^ and mutations or misregulation of ENaC, KV1.1, and Kv1.2 are indicators of poor prognosis in a variety of cancers^49,50^. As epithelial cell extrusion is a fundamental mechanism for epithelial cell turnover, misregulation of extrusion may contribute to disease etiologies arising from mutations in these channels.

Our study identifies early voltage and hydrodynamic regulators as the earliest step controlling homeostatic, crowding-induced cell extrusion, a central driver of epithelial cell death. The channels identified here may be specific for kidney epithelia, yet similar channels expressed in different epithelia^51^ may analogously tune tissues to enable bending and specific threshold densities for activating cell shrinkage and extrusion, depending upon organ function and shape.

## METHODS & MATERIALS

### Cell culture

MDCK II cells (tested for Mycoplasma but not authenticated, gift from K. Matlin, University of Chicago, Chicago, IL) were cultured in Dulbecco’s minimum essential medium (DMEM) high glucose with 10% FBS (Thermofisher) and 100 μg/ml penicillin/streptomycin (Invitrogen) at 5% CO2, 37°C.

### Osmolarity solutions

To test induction of live cell extrusion a range of solution osmolarities were produced. Since MDCKII cells are cultured in DMEM, D-Mannitol (Sigma-aldrich, M4125-1kg) or nuclease-free water (Ambion, AM9937) were added to DMEM to create hyper or hypotonic media, respectively. Initial DMEM osmolarity were measured with a freezing point osmometer (Gonotec, Osmomat 3000), ranged 334-368 mOsm/kg. Solutions were tested bi-weekly and each time a new batch was prepared.

### Immunostaining

Cells were fixed with 4% formaldehyde in PBS at room temperature for 20 min, rinsed three times in PBS, permeabilised for 5 min with PBS containing 0.5% Triton X-100, and blocked in AbDil (PBS 5% BSA). Coverslips were then incubated in primary antibody (in PBS 1% BSA) overnight at 4°C, washed three times with PBS, and incubated in secondary fluorescently-conjugated antibodies. Antibodies used at 1:200 unless otherwise specified: rabbit Piezo1 (Novus, NBP1-78446); mouse S1P (Santa Cruz, CA sc-48356); rabbit KCNA1 (Alomone Labs, APC-161); rabbit KCNA2 (Alomone Labs, APC-010); rabbit LRRC8A (Alomone labs, AAC-001); mouse ZO1 (Invitrogen, 33-9100); rabbit ENaC antibodies SCNNA1 (Invitrogen, PA1-920A), SCNNB1(Invitrogen, PA5-28909), and SCNNG1(Invitrogen PA5-77797). Alexa Fluor 488, 568 and 647 goat anti–mouse and anti–rabbit IgG were used as secondary antibodies (Invitrogen). F-actin was stained using either conjugated 488 or 568 Phalloidin (66μM) at 1:500 and DNA was detected with 1μg/ml DAPI (Themofisher) in all fixed cell experiments.

### Experimental setups and Quantification

#### QPI

Micropatterning MDCK II cells for QPI acquisition relies on having nearby cell-free areas for measurement of cell mass. To achieve this, we grew monolayers on small patterned circles within a dish by adhering a silicone laser cut 100 micromesh disk (Micromesh Array, <u>MMA-0500-100-08-01</u>) on to a non-tissue culture treated 35 mm dish (Ibidi, 81151). The dishes were plasma treated with mesh in place using a chamber (Harrick Plasma, Cleaner PD-32G) applied in a vacuum (Agilent Technologies, IDP-3 Dry Scroll Vacuum pump) for 10 minutes to create cell adherent coating islands with cell free background (Supplemental Figure 1A). Immediately following plasma treatment, the silicone mesh was aseptically removed and MDCKII cells were seeded at 128,000 cell density per well in a microscopy imaging 35 mm dish (Ibidi, 81151) and incubated at 37°C for 6 hours in DMEM. Before filming, excess cells were removed from un-patterned areas by gently washed twice with DMEM and grown another 48 hours.

To image, cells were placed in and on-stage incubator for QPI and islands showing the entire cell island boundary and encompassing empty space were filmed. A minimum of two images of empty space were used for background correction. Images were acquired every 2 minutes for 10 hours at 37°C, 5% CO_2_ and 88% humidity.

Dry mass was then calculated by subtracting the reference images from the island images to correct background. Next background was flattened by subtracting the background average image to flatten the island images. The phase shift was then converted to dry mass using previous established methods^1,2^ in Matlab (version R2022a). Extrusions were quantified from 12 separate islands.

#### Volume quantification

MDCK II cells were plated at 28,000 cell density per well in a microscopy imaging 35 mm dish (Ibidi, 81156) and incubated at 37°C overnight or until 60% confluent. Once 60% confluent, cells were transfected following manufacturer’s protocol with Lipofectamine 3000 (Thermofisher, L3000001) with Cytoplasmic GFP plasmid (addgene, pEGFP-N1 CODE) for 18 hours. Transfection medium was removed, and cells rinsed twice with 1xPBS and incubated in full DMEM until cell to cell junctions were mature (72 hours). To image, cells were stained with Deep Red Cell Mask (Thermofisher, 1.5:1000) for 30 minutes and Hoechst (Invitrogen, 1:1000) in PBS for 10 minutes at 37°C, washed twice with PBS, and then incubated in DMEM media in an enclosed incubated stage at 37°C with 5% CO_2_ (Oko labs) and 0.4 μm Z-slices were captured imaged using a spinning disk microscope (Nikon, Ti2) for up to 10 hours every 10 seconds. Images were analysed using a threshold macro (Nikon Elements AR, 5.41.02) to quantify the volume changes of cells expressing cytoplasmic GFP with cell mask membrane-stained boundaries to highlight extrusions. The volume data was normalized to baseline volume before junctional changes and extrusion in Excel (Microsoft) and graphed and analysed using GraphPad Prism 9.4.1.

#### Serial osmolarity treatment of MDCKII

0.53 x10^5^ cells were grown to confluence (~100 hours) on glass round coverslips 22×55mm (Academy,400-05-21). Epithelial monolayers were incubated with isotonic DMEM for 10 minutes, before treating another 10 minutes in hyper or hypotonic media to induce cell shrinking or swelling, respectively, then replaced in isotonic DMEM for 120 mins. Cells were either filmed (see below) or fixed and stained for quantifying extrusions. Experiment cell density were analysed with bright spots macro in NIS Elements General Analysis (Nikon Elements AR, 5.41.02) using DNA staining to determine cell density per field and phalloidin and DNA to identify extrusions.

#### Live Hypertonic Shock

For live imaging following hypertonic shock, MDCK II cells were plated on an 8 well slide (Ibidi, IB-80801) at a seeding density of 10,000 cells per well and incubated for ~72 hours until monolayers were confluent with mature cell-cell adhesions. Cells were stained with Hoechst (1:1000) in PBS for 10 minutes at 37°C, washed twice with PBS, and incubated in isotonic DMEM media (+/− inhibitors), placed in a microscope stage incubator (37°C, 5% CO_2_, Okolabs), and imaged every 10 seconds for 2.5 hours using a widefield microscope (Nikon, Ti2 specifications below). For live cleaved caspase 3 staining, cells were incubated as per manufactures instructions (1:200, Incucyte^®^ Caspase-3/7 Dyes) before treatment and imaging. All experiments consist of three phases: Baseline: 0-10minutes-cells incubated in isotonic medium (+/− inhibitors, see table below or siRNA knockdown Suppl. Fig 3). Hypertonic Challenge-10-20 minutes: imaging is paused, isotonic medium is syphoned off and replaced with 20% hypertonic medium +/− inhibitors. Imaging promptly restarted to capture shrinkage. Effects on Extrusion-20minutes-end of imaging: imaging is paused and 20% hypertonic medium +/− inhibitors is replaced with isotonic medium +/− inhibitors and timelapse imaging resumed to capture extrusions over the next 2hrs.

#### INHIBITOR TABLE

| **Inhibitor** | **Concentration** | **Target** |
| --- | --- | --- |
| 4AP | 1mM | Voltage-Gated Potassium Channels, e.g. K<sub>v</sub>1.1, K<sub>v</sub>1.2 |
| Azimilide Dihydrochloride (Azi) | 0.5μM | Voltage Gated Potassium Channel K<sub>v</sub>11.1 (hERG) |
| E4031 | 1 μM | Voltage Gated Potassium Channel K<sub>v</sub>11.1 (hERG) |
| Apamin | 10μM | Small Conductance Potassium Channel (SK) |
| Tram 34 | 30μM | Intermediate Conductance Potassium Channel (IK) |
| Iberiotoxin | 100nM | Big Conductance Potassium Channel (BK) |
| GdCl<sub>3</sub> | 10μM | Stretch Activated ion channels |
| GsMTx4 | 10μM | Piezo1, TRPC1, and TRPC6 |
| CaCC inh-A01 | 10μM | Calcium activated Chloride channel |
| CFTR (inh)-172 | 5μM | Cystic Fibrosis Transmembrane Regulator (CFTR) |
| DCPIB | 20μM | SWELL1/LRRC8A |
| Blebbistatin | 100μM | Myosin II Inhibitor |
| Y27632 | 50μM | Rho kinase (ROCK) |
| H1152 | 5nM | Rho kinase (ROCK) |
| SKI-II | 30μM | Sphingosine Kinase |
| SKI V2 | 4μM | Pan S1P receptor |
| JTE-013 | 10μM | S1P<sub>2</sub> receptor |
| zVAD-fmk | 100μM | Pan caspase |

Experiments were then quantified for extrusion rates per 1,000 or 10,000 cells over timelapse video identified by phase microscopy or % shrinkage by Lightning Assays described below.

#### siRNA knock down

4-siRNA smart pools (Horizon Discovery, L-006210-00-0010 (Kv1.1), L-006212-00-0010 (Kv1.2), L-026211-01-0010 (SWELL1), L-006504-00-0010 ENaCα, L-006505-00-0010 (ENaCβ) or L-006507-01-0010 (ENaCγ) or D-001810-01-20 (non-targeting control)) were prepared to in DNase RNase free water to a 100 μM stock. MDCK II cells were seeded in tandem one set on a 6 well plate (Thermofisher, 140675) at 28,000 cell density (for qPCR analysis) and the other on a 8 well slide (Ibidi, IB-80801) at 10,000 cell density (for live cell imaging) or on coverslips seeded with 53,000 MDCK II cells per well on a 24 well plate (Thermofisher, 142475) for extrusion and buckling quantification (described below), and grown overnight until 60% confluent. Cells were then transfected with RNAi Max kit (Thermo Fisher, 13778150) and 1μM siRNA for 24h before replacing with fresh DMEM for 48h. Cell knockdowns plated for RT-qPCR analyses were lysed for RNA extraction using RNAeasy kit (Qiagen, 74104), following the manufacturer’s instructions. One μg of RNA was purified with 1 μL of 10x DNAse I Reaction Buffer, 1 μL of DNAse I Amplification and RNase-free water in a final volume of 10 μL. The samples were incubated for 10 min at 37°C then enzyme was deactivated with 1 μL of 0.5M EDTA for 10 min at 75°C. Samples were stored at −20°C. or directly processed by RT-qPCR using Brilliant III Ultra-Fast SYBR Green QRT-PCR Master Mix (Agilent Technologies), using primers designed with SnapGene^®^ (version 6.1.1) below and produced by Sigma Aldrich (see Supplemental Fig 3B for each). Reactions were analysed with ViiA 7 Real-Time PCR System (Thermofisher) using the following cycle conditions: 50°C for 10 minutes, 95°C for 3 minutes, followed by 40 cycles at 95°C for 15 seconds and 60°C for 30 seconds. Results were normalized to Glyceraldehyde 3-phosphate dehydrogenase (GAPDH) expression and graphed and statistically analysed with GraphPad Prism 9.4.1. Extrusion rates were quantified per 1,000 cells using timelapse phase microscopy identified and % shrinkage using Lightning Assay, described below.

##### TABLE OF SEQUENCES

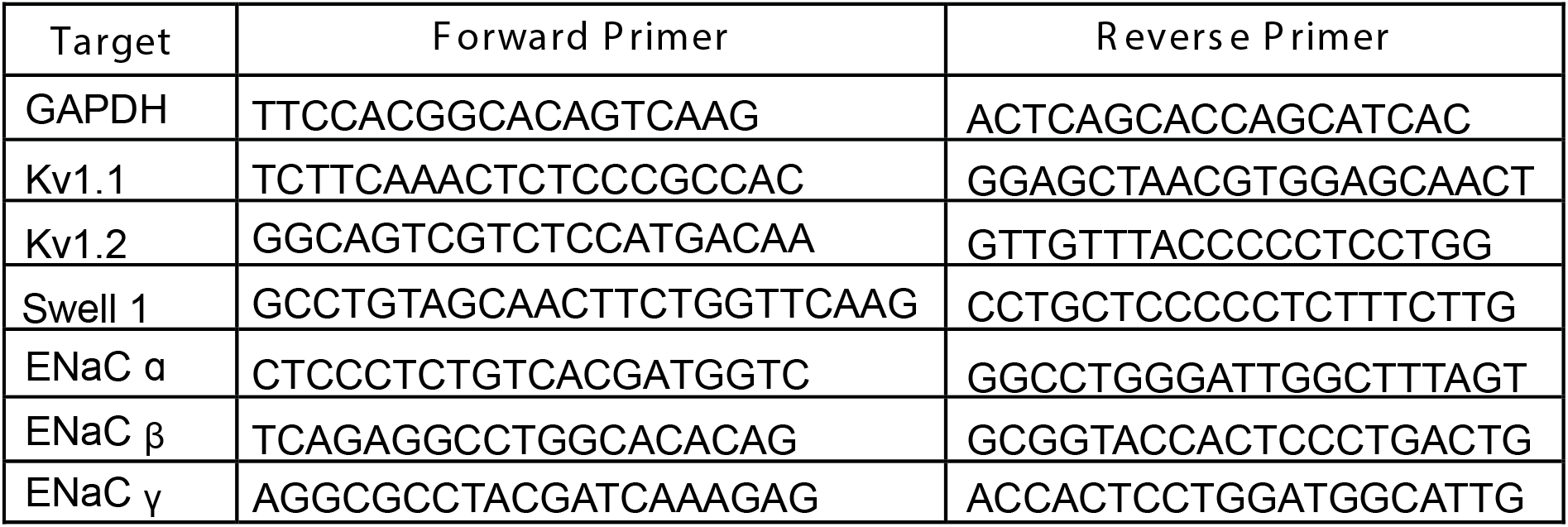

#### The Lightning Assay

To expedite analysis of cell shrinkage following modulation of different channels, we used the ‘lightning assay’. To do this, areas of interest were cropped out of larger phase microscopy timelapse movies. Junction intensity changes were collected by applying a simple threshold macro based on white light detection before, during, and briefly after extrusion. The threshold was set to capture the area changes around the cells in the frames before homeostatic or experimental cell shrinkage. The same threshold was applied to all frames of movie until completion of cell extrusion. This same method was used for both single cell and whole regions of crowding in an 85 μm^2^ (400 by 400-pixels) area. Data were then normalized in Microsoft Excel (version 16.67) using an average of 10 frames before lightning and analysing peak percentage change, and then graphed and statistically analysed using GraphPad Prism 9.4.1.

#### Buckling

MDCK II cells were seeded at 0.53 x10^5^ on glass round coverslips 22×55mm (Academy, 400-05-21) for fixed imaging or seeded at 1.28 x10^5^ cells per well in a 35 mm dish for live imaging. After seeding cells were grown overnight, and transfected with either non-targeting control, KV1.1, KV1.2, SWELL1, ENaC-α, β, or γ siRNA, as described above. After transfection coverslip plated epithelial cells were incubated for cover 48 hours at 37°C, then coverslips were fixed and immuno-stained. siRNA-treated MDCK II cells were imaged using a spinning disk confocal microscope with 0.4μm thick sections and a maximum height of 32μm.

After transfection of epithelial cells, plated on 35mm dishes, were incubated 24 hours then live imaging was completed using the widefield microscope. Phase images were captured every 10 minutes for 10 or more hours. For both live and fixed buckle analysis, buckles were scored as mounds with more than 3 connected cells within each field and analysed in Graph Pad Prism 9.4.1.

#### Depolarization

MDCK II cells were seeded at 128,000 cells per well in a 35 mm dish and grown ~72 hours to confluence and junctional maturity. Once mature, monolayers were stained with DiBAC_4_(3) following protocol entitled “Tracking transmembrane voltage using DiBAC_4_(3) fluorescent dye (PDF)” at: https://ase.tufts.edu/biology/labs/levin/resources/protocols.htm). Monolayers were then treated with DMSO (vehicle), 4-AP, or amiloride and imaged (phase and GFP settings) every 10 seconds for a minimum of 2.5h. The images were analysed using Nikon Elements AR, 5.41.02, using a region of interest (ROI) over any cell that was maintained or extruded. ROI mean intensity of DiBAC_4_(3) in each cell over time was normalized in Excel using 10 baseline frames before shrink and depolarization over time was graphed in GraphPad Prism 9.4.1. For neighbour voltage differentials, the cells immediately in contact with an extruding cell were classified as brighter, equal, or darker than the extruding cell, based on DiBAC_4_(3) intensities. Cell counts where then plotted and analysed in Graph Pad Prism 9.4.1

### Microscopy

#### QPI

Time-lapse QPI and brightfield images were collected on an Olympus IX83 inverted microscope (Olympus Corp) using a 40x NA 0.75 objective. Samples were illuminated using red LED light (623nm, Thorlabs, DC2200) for 120ms exposure a QWLSI wavefront sensing camera-(Phasics SID4-4MP), driven by Micro Manager opensource microscopy software. Samples were incubated with a stage-top incubator (Okolabs) set at 37°C temperature with 5% CO_2_ gas and 95% humidity.

#### Widefield

Timelapse phase and fluorescence images were captured on Nikon Eclipse Ti2 using a Plan Fluor 20x Ph1 DLL NA=0.50 objective with a Photometrics Iris 15 16-bit camera and a Cool LED pE-4000 lamp driven by NIS Elements (Nikon, version 5.30.02).

#### Spinning disk

Images were captured on a Nikon Eclipse Ti2 using 20x, 40x plan fluor 0.75 air or 60X or 100X plan fluor 1.40 oil objectives with an iXon 888 Andor 16-bit camera, a Yokogawa CSU-W1 confocal spinning disk unit and Toptica photonics laser driven by NIS Elements (Nikon, 5.21.03). Cell staining with phalloidin and Hoechst were quantified for extrusions per 1,000 or 10,000 cells using Nikon Elements Software.

#### Statistical analysis

For statistical analysis, all experiments were repeated on at least three separate days to capture variation in the biological replicates. Data were analysed using statistical software GraphPad Prism 9.4.1 to measure normality by Shapiro-Wilk test and significance by unpaired t-tests, Two-Way ANOVA, or One-way ANOVA with Dunnett’s multiple comparisons, Tukey’s or Sidak’s corrections as described in figure legends. To reduce bias, we imaged random fields within the middle or crowded areas of glass coverslips for quantification or the center of an 8 well dish (Ibidi 80806) for live cell shrink experiments. We excluded low-density epithelia (fields with less than 2000 cells) as these regions are not crowded enough to elicit extrusion. Graphs were generated using GraphPad Prism 9.4.1. Figure layouts and models were created in Adobe Illustrator (26.3.1).

1. Pradeep, S. & Zangle, T A. Quantitative phase velocimetry measures bulk intracellular transport of cell mass during the cell cycle. Sci Rep 12, 6074 (2022). https://doi.org:10.1038/s41598-022-10000-w
2. Zangle, T A. & Teitell, M. A. Live-cell mass profiling: an emerging approach in quantitative biophysics. Nat Methods 11, 1221-1228 (2014). https://doi.org:10.1038/nmeth.3175

## ACKNOWLEDGEMENTS

We especially thank Michael Redd for his typical in-depth questioning that made us revise models and improve the manuscript and also to Alexandra Berr, Dustin Bagley, John Fadul, Tobias Russell, Faith Fore, Laura Akin-tche, and Lily Gates for helpful comments on our manuscript. J.R. is the recipient of a National Institute of Health R01GM102169, a Howard Hughes Faculty Scholar Award (55108560), a Cancer Research UK Programme Grant (DRCNPG-May21\100007), an Academy of Medical Sciences Professorship (APR2\1007), and a Wellcome Trust Investigator Award (221908/Z/20/Z) C.P.-P. is the recipient of a Long-Term Fellowship (LT000654/2019-L) from the Human Frontier Science Program organization and a Marie Skłodowska-Curie Fellowship (898067) from the European Union’s Horizon 2020 research and innovation program. This work was supported by the Office of the Assistant Secretary of Defense for Health Affairs through the Breast Cancer Research Program under Award Number W81XWH1910065 (TAZ).

## Contributions

J. R. and S. J. M., and designed experiments, interpreted, analyzed data, and wrote the manuscript. C. P. P. co-designed experiments and interpreted data. T. A. Z. designed QPI experiments and helped with their interpretations. All authors edited the manuscript.

## Supplemental Videos

**S Video 1:** MDCKII dry mass changes before extrusion. Images taken every 2 minutes using QPI adapted technique shown in parula color map.

**S Video 2:** Phase timelapse (hh:mm) of MDCKII cell show junctional lightning before extrusion.

**S Video 3:** Phase and cleaved caspase 3 (green) timelapse of apoptotic MDCKII cell that does not show junctional lightning before extruding.

**S Video 4:** Phase microscopy timelapse (hh:mm) of MDCKII monolayer in isotonic media.

**S Video 5:** Phase microscopy timelapse (hh:mm) of MDCKII cells after ten-minute incubation in 20% hypertonic media undergo increased rate of extrusion.

**S Video 6:** Phase and DiBAC_4_(3) (green) timelapse (mm:ss) showing depolarization of MDCKs before extrusion.

**S Video 7:** Phase and DiBAC_4_(3) (green) timelapse (mm:ss) showing differential depolarization as MDCKs crowd (red arrows) and extrude (white arrows). Scale bar (white) 25mm.

**S Video 8:** Representative phase timelapse (mm:ss) of MDCKII monolayers where Swell1 is knocked down, which buckle over time, with scale bar 25mm.

## Notes

### Competing Interest Statement

The authors have declared no competing interest.

